# Gene PointNet for Tumor Classification

**DOI:** 10.1101/2024.06.02.597020

**Authors:** Hao Lu, Mostafa Rezapour, Haseebullah Baha, Muhammad Khalid Khan Niazi, Aarthi Narayanan, Metin Nafi Gurcan

## Abstract

The rising incidence of cancer underscores the imperative for innovative diagnostic and prognostic methodologies. This study delves into the potential of RNA-Seq gene expression data to enhance cancer classification accuracy. Introducing a pioneering approach, we model gene expression data as point clouds, capitalizing on the data’s intrinsic properties to bolster classification performance. Utilizing PointNet, a typical technique for processing point cloud data, as our framework’s cornerstone, we incorporate inductive biases pertinent to gene expression and pathways. This integration markedly elevates model efficacy, culminating in developing an end-to-end deep learning classifier with an accuracy rate surpassing 99%. Our findings not only illuminate the capabilities of AI-driven models in the realm of oncology but also highlight the criticality of acknowledging biological dataset nuances in model design. This research provides insights into application of deep learning in medical science, setting the stage for further innovation in cancer classification through sophisticated biological data analysis. The source code for our study is accessible at: https://github.com/cialab/GPNet.

## 1. Introduction

Cancer ranks among the most lethal diseases, and its growing incidence underscores the urgency for early detection and accurate diagnosis and prognosis [1-3]. Scientists and researchers are continually seeking innovative methods to combat this challenge. Gene expression data, like RNA-Seq data, have been proven to be a powerful approach to diagnosing and prognosing specific cancer types [4-6]. With the increase in the volume of gene expression data, AI-driven deep learning models have demonstrated better performance in key feature extraction and gene profile classification compared to traditional analysis techniques and shallow models.

For RNA-Seq data analysis, most current studies [7-9] use Fully Connected networks (FCNs). According to the universal approximation theorem [10], the FCNs can approximate any continuous function. This suggests that for data with unknown structure and distribution, FCNs are often the first choice. However, FCNs treat each input feature as potentially important on its own without necessarily considering its relationship or proximity to other features. This means the network does not assume, for instance, that features in close proximity to each other in the input vector are more potentially related than features further apart. Nonetheless, many studies [11, 12] have shown that genes do not act in isolation but are part of larger signaling and metabolic pathways. Genes involved in the same biological pathways or cellular processes often exhibit co-expression patterns, meaning their expression levels are correlated across different samples or conditions. Meanwhile, previous studies demonstrated that the correct assumption that a learning algorithm makes about the data, which guides the learning process and helps the model generalize well from training data to unseen data, can expedite model convergence, enhance predictive accuracy, and reduce the model’s dependency on large data volumes [13-15]. These assumptions are referred to as *inductive biases*.

A classic example of introducing good inductive biases is in the design of Convolutional Neural Networks (CNN). CNNs inherently assume that the data has a hierarchical spatial structure, only a small subset of the input data (pixels in the case of images) is relevant to a particular computation, and the patterns to be detected are the same throughout the input space. These inductive biases, specifically designed for the characteristics of image data, have led to tremendous success for CNNs in the field of image analysis [16, 17]. This insight has led us to explore new methodologies in our approach to intentionally introduce inductive biases suitable for the properties of gene expression files.

This paper models gene expression data as a point cloud data structure and employs PointNet [18], a standard approach, as the backbone for processing gene expression data represented as point clouds. This innovative method allows us to leverage the spatial relationships within the data, offering a fresh perspective on gene expression analysis. PointNet is also amenable to us introducing inductive biases. Another innovative approach is to use a classifier combined with gene pathway knowledge. By integrating these elements, our study not only advances the technical capabilities of deep learning models but also enriches our understanding of the biological underpinnings of cancer. This approach underscores the significance of considering the intrinsic properties of data in deep learning research, particularly in the context of complex biological data sets like gene expression profiles. We expect this article to inspire other researchers studying gene expression data to introduce more reasonable inductive biases (much like the early works on CNN) and ultimately find a network structure suitable for gene expression data.

The interpretability of these models is as important as their development and performance. We can interpret models using explainable methods, such as Class Activation Mapping (CAM) [19, 20]. These methods enable an in-depth examination of the algorithm’s attention allocation to individual genes during the cancer classification process, allowing for the inference of associations between specific genes and cancer types. This not only advances our scientific knowledge but also opens the door to more personalized and precise cancer treatments, highlighting the crucial role of AI in modern medicine. These methods enable an in-depth examination of the algorithm’s attention allocation to individual genes during the cancer classification process, allowing for the inference of associations between specific genes and cancer types.

Building upon these capabilities, our study introduces a novel framework, GenePointNet (GPNet), designed to harness the power of gene expression data for cancer classification through an innovative application of point cloud data structures and deep learning techniques. Our approach operationalizes the gene point cloud data structure concept through a structured methodology that unfolds in four critical steps:
1. **Data Preprocessing:** The gene expression data undergoes a rigorous preprocessing phase to reduce noise and filter out irrelevant information. This ensures that the subsequent analysis is based on clean and reliable data (Section 3.1).
2. **Point Cloud Generation:** Post-preprocessing, the refined gene expression data is transformed into a point cloud format, each point representing a gene. This representation captures the spatial and relational dynamics of gene expressions in a novel manner (explained in Section 3.2.1).
3. **Deep Learning Model:** Leveraging the point cloud data, we employ a sophisticated deep learning model adept at classifying the points based on their characteristics and underlying biological knowledge. The model is meticulously trained on a comprehensive dataset comprising labeled cancer samples, facilitating a robust learning mechanism (outlined in Section 3.2.2).
4. **Classification:** The culmination of this process is the model’s ability to classify new cancer samples accurately, marking a significant advancement in precision oncology (elaborated in Section 3.2.3).

The contributions of this paper are manifold and underscore the innovative aspects of our research:

1. We introduce the modeling of gene expression data as a point cloud, deliberately integrating inductive biases pertinent to gene expressions and gene pathways into our deep-learning model, GPNet. This novel approach not only enhances the accuracy of cancer classification but also aligns with the intrinsic properties of gene expression data, marking an advancement in the field.
2. Through careful development and optimization, we introduce an end-to-end deep learning classifier specifically tailored for cancer classification. Our model demonstrates a remarkable accuracy rate exceeding 99%.
3. Additionally, our study identifies the top three genes identified by our model, enabling us to explore their relationship with tumor processes. These insights not only contributes to our understanding of cancer mechanisms but also paves the way for the discovery of more effective cancer biomarkers, offering promising avenues for future research.

## 2. Related works

Since the early development of gene expression profiling technology, it has been a valuable tool for cancer classification. Golub et al. [4] demonstrated the potential of microarray expression profiles to distinguish between types of leukemia, showcasing the clinical utility of gene expression signatures despite the challenges in interpreting them in biological terms [21]. The advent of deep learning technology has seen its application extend into tumor classification, with research efforts diversifying into four main categories:

- *Convolutional Neural Network (CNN) Models for Classification:* Shon et al. [22] utilized PCA to reduce the dimensionality of gene expression data, reshaping it into 2D HeatMaps for CNN-based classification. Similarly, Mostavi et al. [23] investigated using 1D convolutional networks for classifying original data and 2D CNNs for data reshaped into 2D matrices.
- *Artificial Neural Network (ANN) Models for Classification:* Dwivedi [7] applied ANNs to classify leukemia subtypes, demonstrating their superiority over traditional ML methods. Urda et al. [8] also employed ANNs for tumor classification, preceded by dimensionality reduction techniques like t-tests or lasso regression to filter out highly correlated genes.
- *Autoencoders for Dimensionality Reduction:* Autoencoders can further advance dimensionality reduction. Xiao et al. [9] utilized a stacked sparse autoencoder for dimensionality reduction before classification with ANNs. Teixeira et al. [24] implemented a stacked denoising autoencoder for the same purpose, comparing the efficacy of autoencoders against linear (PCA) and nonlinear (KPCA) dimensionality reduction methods.
- *Transformer Models in Single-Cell RNA-Seq:* The maturation of single-cell RNA-Seq technology and the adaptability of transformer models have led to new research avenues [25-28]. TOSICA [20] represents the state-of-the-art in cell-type prediction without relying on self-supervised pre-training. This paper contrasts a fine-tuned pre-trained TOSICA model against our model on smaller datasets to highlight our model’s efficacy.

This study introduces a novel approach by modeling gene profile data as point clouds and employing a lightweight PointNet for feature extraction. PointNet [18], a pioneering 3D point cloud analysis model, processes point clouds directly while maintaining permutation invariance, which is essential for their irregular format. Its application to tasks such as classification and segmentation has demonstrated robustness against noise and occlusion, establishing it as a foundational model in 3D computer vision and inspiring further developments in point cloud processing.

## 3. Materials and Methods

### 3.1. Data

This study utilized publicly accessible RNA-Seq gene expression datasets from the Genomic Data Commons (GDC) [29], which are available for download at no cost through their official website. We curated a comprehensive dataset by combining gene expression data from four distinct projects: Acute Myeloid Leukemia (AML), Breast Invasive Carcinoma (BRCA), Colon Adenocarcinoma (COAD), and Kidney Renal Papillary Cell Carcinoma (KIRP). This compiled dataset encompasses 4916 samples across six tumor cell types and 594 samples from normal cells adjacent to the tumors, yielding a total dataset size of 60660 genes per sample. **Table 1** provides a detailed overview of the RNA-Seq dataset composition, including the classification of samples into normal (N) and tumor (T) categories, along with the number of samples for each class:

**Table 1.**
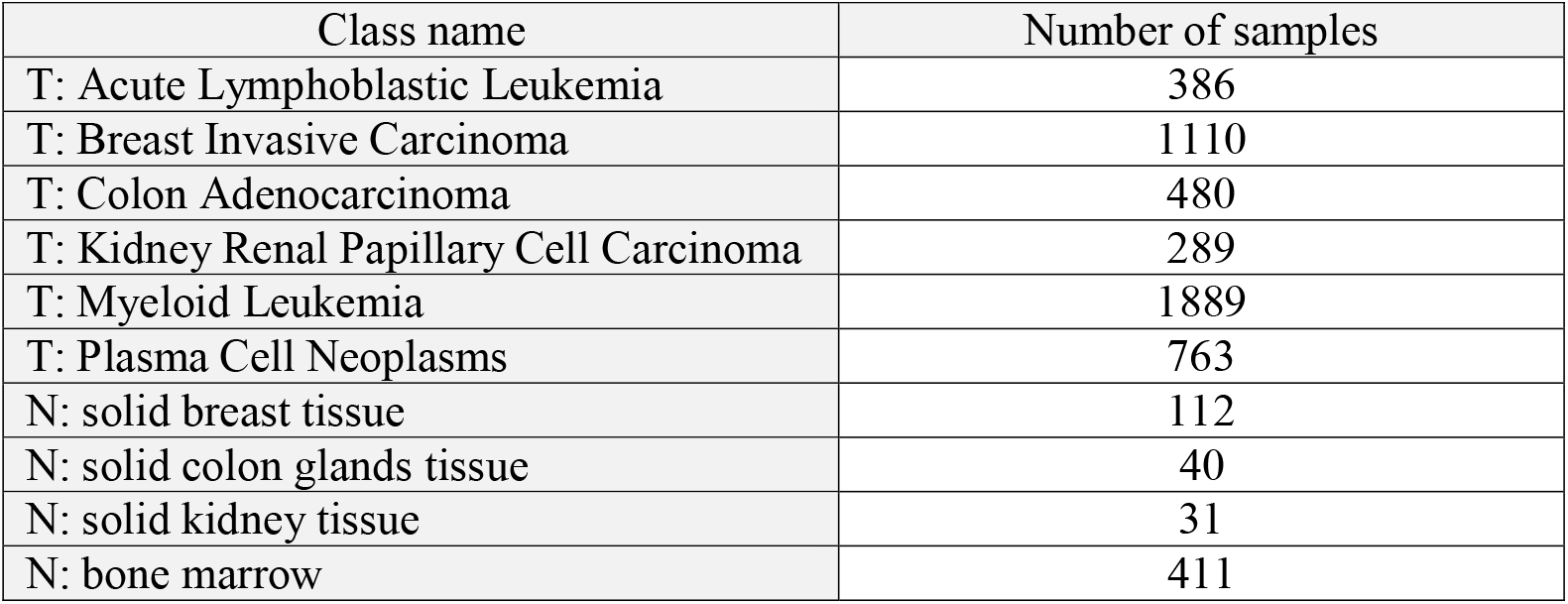
Information of our built RNA-Seq datasets: normal samples (N), tumor samples (T), number of samples (#sample)

| Class name | Number of samples |
| --- | --- |
| T: Acute Lymphoblastic Leukemia | 386 |
| T: Breast Invasive Carcinoma | 1110 |
| T: Colon Adenocarcinoma | 480 |
| T: Kidney Renal Papillary Cell Carcinoma | 289 |
| T: Myeloid Leukemia | 1889 |
| T: Plasma Cell Neoplasms | 763 |
| N: solid breast tissue | 112 |
| N: solid colon glands tissue | 40 |
| N: solid kidney tissue | 31 |
| N: bone marrow | 411 |

This heterogeneous dataset provides a robust foundation for applying our proposed GenePointNet (GPNet) framework to the task of cancer classification via RNA-Seq data.

### 3.2. Structure of Gene PointNet

This study introduces an innovative approach to representing RNA gene expression data, conceptualizing it as the amount of gene expression akin to how light intensity in a pixel represents an image. This analogy allows us to draw parallels between gene expression profiles and digital images, where the expression level of a gene is akin to a pixel’s light intensity. In images, this intensity is closely related to the pixel’s immediate neighbors, mirroring how, in gene expression data, a gene’s expression level is closely related to that of functionally similar genes, such as those within the same pathway.

Unlike the uniform, equidistant relationships between neighboring pixels in a 2D image, gene expression profiles feature a variable number of connections reflecting the complex interplay of gene functions. This complexity is depicted in **Figure 1**, illustrating the transition from 1D and 2D data structures to a high-dimensional point cloud representation. Each gene (point) interacts with multiple neighbors at varying distances in this high-dimensional space, as shown in **Figure 1(c)**. So we map the gene expression matrix to a point cloud. The main idea of this paper, as shown in **Figure 1(d)**, is that we aim to cluster the points corresponding to genes that are functionally related and relevant to tumor categories through an end-to-end training process. Then, by differentiating the expression counts of these genes, we distinguish between different tumor classes. It is crucial to note that, distinct from traditional point cloud data, which can vary in shape, the spatial configuration of genes in a gene point cloud remains constant across individuals of the same species, with variations manifesting in the expression levels rather than the positions of genes.

**Figure 1.**
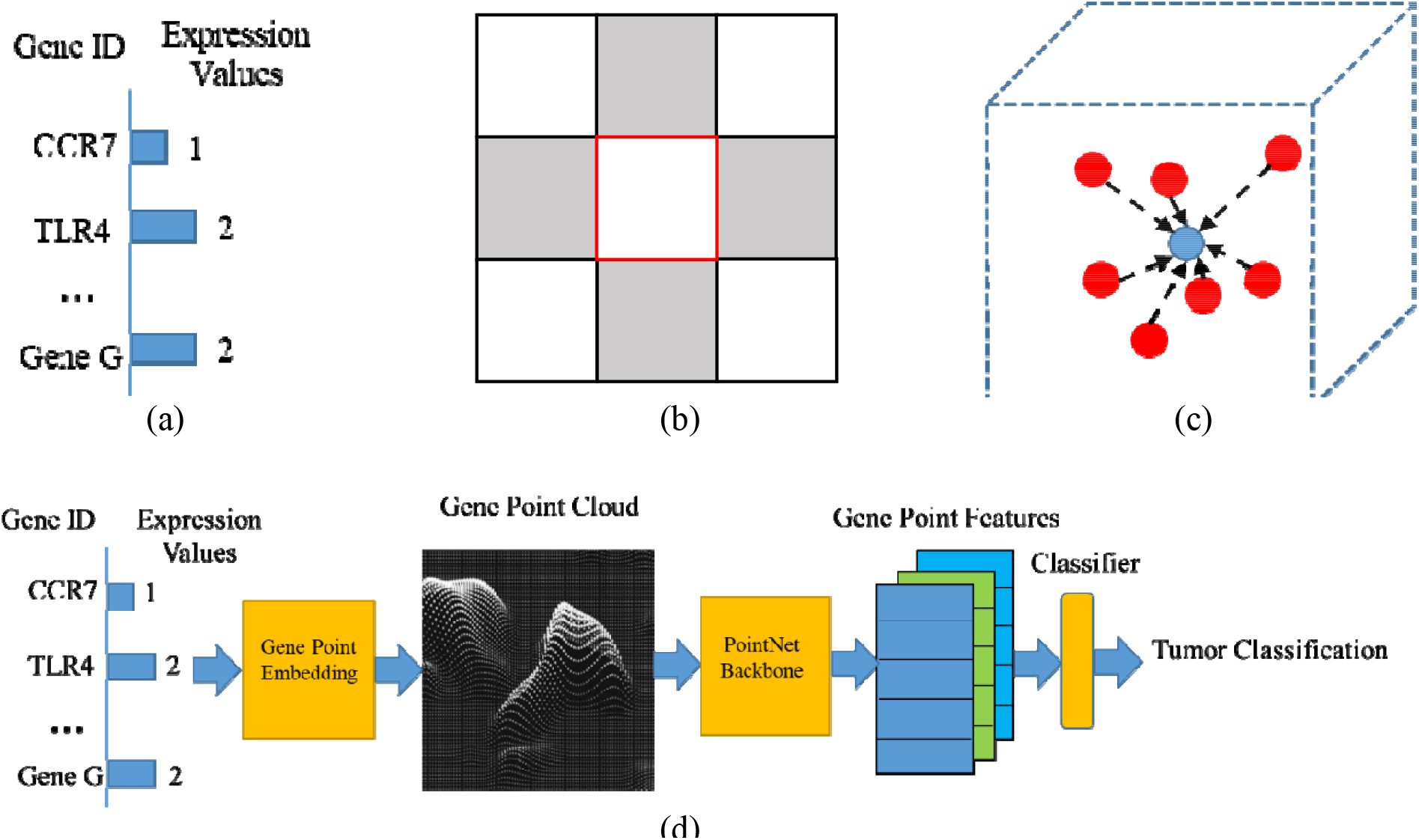
(a) Gene expression values that can be represented by a 1D data structure. (b) 2D image data structure, where each pixel has 8 neighbor pixels. (c) High-dimension point cloud data structure, where each point has a multitude of heterogeneous neighbors. (d) Schematic diagram of Gene PointNet end-to-end training process.

To process this complex data, we apply PointNet, as proposed by Qi et al. [18], designed to handle point cloud data efficiently. PointNet is a neural network architecture designed to process and classify point clouds consisting of sets of 3D points arranged spatially. Point clouds find common applications in computer vision, robotics, and graphics for tasks like 3D object recognition and scene understanding. The main innovation of PointNet lies in its efficient neural network architecture that can directly process point clouds without the need for any pre-processing or voxelization. This is accomplished by creating a symmetric function that can aggregate information from all points in the cloud while maintaining the input’s permutation invariance.

PointNet comprises two key components: the feature extraction module and the classification module. The feature extraction module utilizes a series of multi-layer perceptrons (MLPs) on each point within the point cloud, followed by a max pooling operation to generate a global feature vector. The classification module then applies another set of MLPs to the global feature vector to predict the class label of the point cloud. We enhance PointNet by substituting its global max pooling layer with a Knowledge-based MLP, inspired by the work on TOSICA [25]. This modification integrates biological insights on gene pathways into the model, facilitating a more informed prediction that accounts for each gene expression’s impact. This adjustment embeds human knowledge into the neural network and streamlines PointNet’s architecture, making it better suited for smaller datasets. **Figure 2(a)** provides a detailed schematic representation of the architecture of our Gene PointNet (GPNet), specifically designed for processing and classifying gene expression data as a high-dimensional point cloud. The architecture is segmented into distinct components, each contributing to the overall functionality of GPNet:

**Figure 2.**
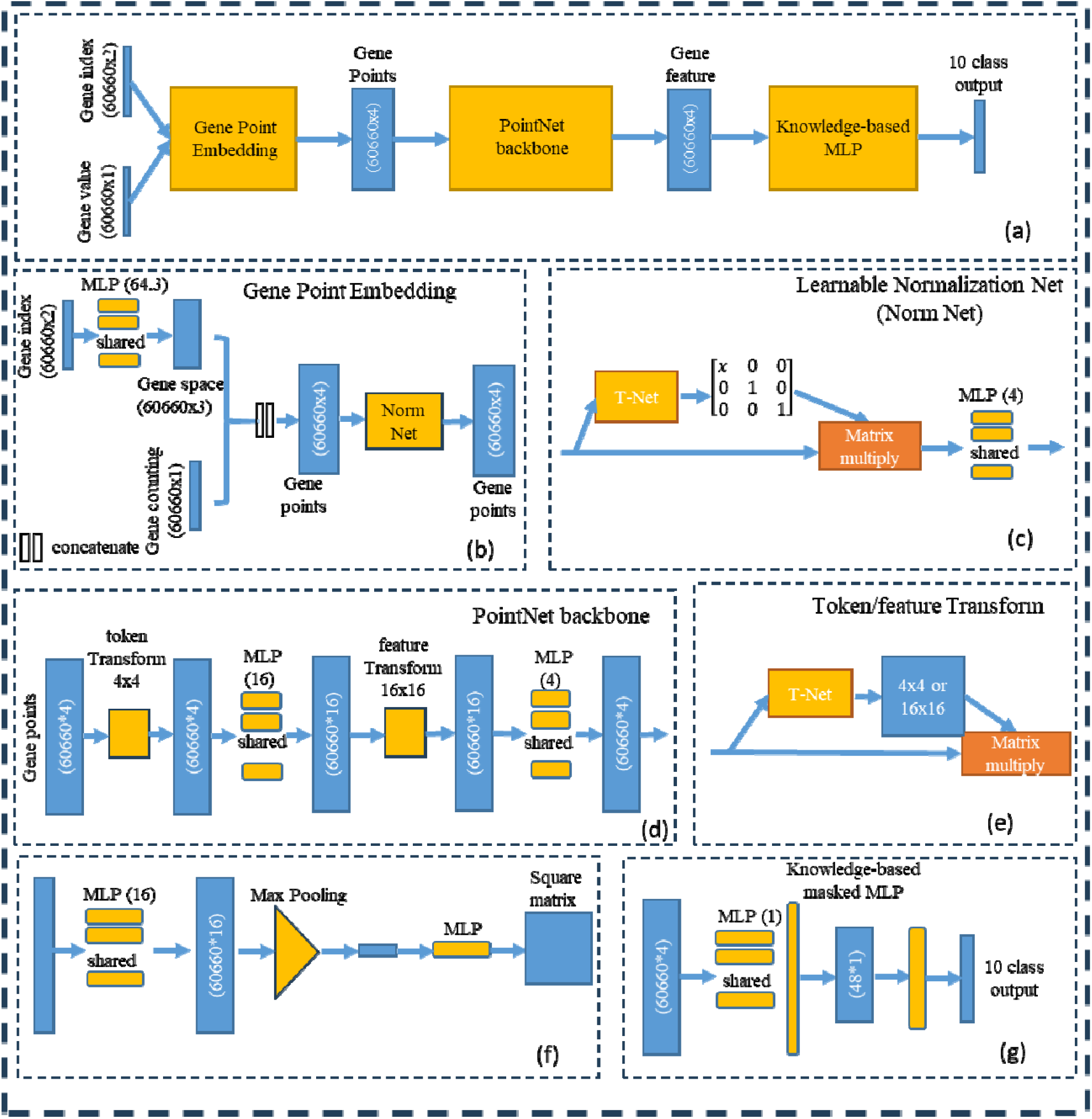
Overall structure for our GPNet (a), (b) shows the gene embedding network, (c) shows the detail structure of learnable normalization network, (d) shows the PointNet[18] backbone for feature extraction of each gene, (e) shows the input/feature transform network in PointNet, (f) shows the details of T-Net in PointNet, and (g) shows the Knowledge-based MLP mentioned in TOSICA[25].

1. **Gene Point Cloud Embedding:** The foundation of GPNet begins with transforming gene expression data into a high-dimensional point cloud format, as illustrated in **Figure 2(b)** and elaborated in Section 2.2.1. This transformative process incorporates gene embedding to represent each gene as a point in high-dimensional space, inputting expression values that capture the gene’s expression level, and a learnable normalization network to standardize these values, ensuring consistent processing across all samples. This normalization process is graphically represented in **Figure 2(c)**, highlighting the initial preparation of gene expression data for subsequent analysis.
2. **PointNet Backbone:** Central to our architecture is a customized, lightweight version of PointNet, responsible for extracting features from the point cloud data. This adaptation, presented in **Figure 2(d)** and discussed in Section 2.2.2, efficiently processes each gene point, extracting crucial features for classification while maintaining a manageable computational footprint. Modifications to the original PointNet design include parameter reduction and structural adjustments, and a lightweight T-Net, which is depicted in **Figure 2(f)**. These changes ensure the model’s capacity to encode gene structure information accurately, fostering an understanding of gene relationships and functional groupings within the point cloud.
3. **Knowledge-based MLP Classifier:** The culmination of the GPNet process involves a knowledge-based MLP classifier, as shown in **Figure 2(g)** and described in Section 2.2.3. This classifier integrates the extracted point features with biological knowledge, employing a convolutional MLP and a knowledge-based masked mapping similar to the approach used in TOSICA [25]. A linear classifier then uses this rich, contextualized feature set to perform the final cancer classification, leveraging both the intrinsic data patterns and external biological insights.
4. **Loss Function and Model Explainability:** Essential to the training and evaluation of GPNet are the considerations of the loss function and model explainability, addressed in Sections 2.2.4 and 2.2.5, respectively. The architecture employs a weighted cross-entropy loss to manage dataset imbalances, depicted in **Figure 2(e)**, and orthogonal losses to preserve the structural integrity of gene embeddings. Moreover, implementing a grad-CAM method, as part of our model’s explainability strategy, allows for the visualization of the model’s focus within the gene point cloud, offering insights into the decision-making process.

GPNet stands as a comprehensive framework for the advanced analysis of gene expression data, embodying a novel approach to cancer diagnosis through the lens of AI and computational biology. By meticulously mapping each step of the process and integrating both data-driven and knowledge-based elements, GPNet promises to enhance our understanding and classification of cancer at the genetic level. Throughout this paper, uppercase bold letters (e.g., **X**) denote matrices, lowercase bold letters (e.g., ***t***_*j*_) denote vectors, and non-bolded letters (e.g., *a*_*j*_) denote scalars.

#### 2.2.1 Gene Point Cloud Embedding

In this subsection, we aim to transform gene profiling data into a gene point cloud. Previously, we discussed how each point’s “color” reflects its expression level, with its spatial position determined by the gene ID. This subsection outlines our strategy for embedding gene IDs, preprocessing expression values, and utilizing a learnable normalization layer to mitigate batch effects in gene expression data.

The RNA-Seq data is formatted as a matrix,**X** ∈*Z*^*NxG*^, where N represents the number of samples, G represents the number of genes, and each element X_*ij*_ ∈Z^+^ corresponds to the RNA read count of gene *j* = 1,2,…, *G* in sample *i* = 1,2,3,…,*N* Specifically, this value represents the abundance of gene *i* (where *i* ∈ 0,1,…*G*) in sample *j* (where *j* ∈ 0,1,…*N*). Throughout this paper, we refer to this matrix as the raw counting matrix.. (where) in sample Inputs to GPNet are twofold: the gene identifier and the expression values, as depicted in **Figure 2(b)**. These components are integrated into the gene point cloud through the following methodology:

##### Gene embedding

In our approach, each gene is represented as a point within the gene point cloud, drawing parallels to how tokens function in natural language generation (NLG). This conceptual similarity allows us to employ a strategy similar to previous studies [26, 30] where the gene name serves as the token. Consequently, we assign to each gene,*g*_*i*_, a unique integer pair as its identifier, 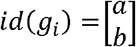 where *a,b* ∈ *N*, among a comprehensive set of 60660 genes. This enumeration integrates all human genes into a unified vocabulary by amalgamating all identified genes across various studies. Therefore, the gene tokens for each sample *j* are represented as a vector ***t***_*j*_ ∈ N^2×60660^, constructed as

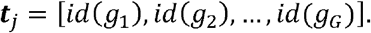

Subsequently, these identifiers are translated into gene points through a convolutional embedding layer, as depicted in **Figure 2(b)**. The resulting embedded gene points, denoted as *mb*_*g*_ **(*t***_*j*_) ∈ N^2×60660^, illustrate how the embedding layer groups related genes while distinguishing unrelated ones.

##### Processing Expression Values

Traditionally, preprocessing of the gene expression matrix **X** involves steps like transcripts per million (TPM) normalization and log1p transformation to approximate a Gaussian distribution of the input data [26]. However, in our approach, we have developed a learnable normalization network specifically designed to automatically normalize the gene expression data. Hence, we do not need the complex traditional preprocessing approaches. Considering that deep learning models converge more easily with zero-mean, Gaussianized data, we only perform simple z-score normalization [31] on the data:

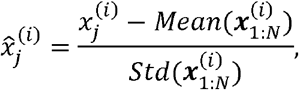

where *i* = 1,2,…, *G* denotes the gene index, and *j* = 1,2,…, *N* denotes the sample index. 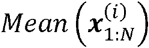 and 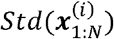 represent the mean and stander derivation for gene *j* across *N* samples, respectively. The resultant vector of normalized expression values for each sample *j* is expressed as:

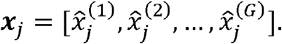

These expression values are then combined with the gene embeddings to form the input for the learnable normalization network:

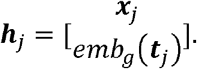

##### Learnable Normalization Network

Normalizing gene profiling data is a crucial step in the analysis of gene expression, applicable to data derived from microarrays, RNA-seq, and other high-throughput methodologies. The necessity for normalization arises from the need to mitigate various sources of variation and bias that can skew gene expression measurements, ensuring that the observed expression differences accurately reflect biological variations rather than technical discrepancies. In our approach, we employ a convolutional structure similar to the T-Net found in PointNet [18] to derive normalization parameters for expression values. This setup comprises:

1. A Convolutional MLP Layer with a ReLU activation function to initiate the normalization process.
2. A Global Max Pooling Layer to aggregate features across the entire dataset, focusing on the most significant signals.
3. An MLP Layer designed to map ***h***_*j*_ to a singular normalization parameter *a*_*j*_. This parameter is then applied to the expression values ***x***_*j*_ to produce normalized expression data 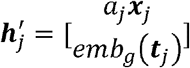. Following normalization, ***h***′*j* is processed through another convolutional MLP layer to generate the final gene point representation:

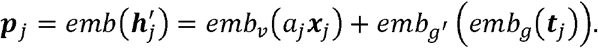

#### 2.2.2 Gene PointNet Backbone

Our model adopts the foundational architecture of the original PointNet but with a reduced parameter footprint, as depicted in **Figure 2(d)**. This streamlined version includes:

- A Convolutional MLP Layer: Serves as the initial layer to process the gene points.
- Two Transformation Networks (T-Nets): One for token transformation (termed input transform in the original PointNet [18]) and another for feature transformation. These are critical for aligning and normalizing the input data into a standardized format conducive to further processing.

However, to tailor this architecture to our specific needs, we have modified it considerably. The modifications of T-Net, detailed in **Figure 2(f)**, focus on preserving the gene point cloud’s structural integrity by introducing orthogonal constraints on both the token and feature transformations. This ensures that the PointNet backbone does not alter the inherent spatial relationships within the gene point cloud, thereby maintaining the encoded gene structure information within *emb*_*g*_(***t***_*j*_) and facilitating the grouping of related genes while distinguishing unrelated ones.

The mathematical representation of this process is as follows:

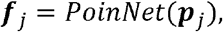

where ***f***_*j*_ ∈ R^4×60660^ represents the output feature matrix for sample *j*, derived from processing the gene point representation ***p***_*j*_ through the modified PointNet architecture. This feature matrix captures the essential characteristics of each gene within the sample, reflecting both its expression levels and its relative positioning and relationships within the high-dimensional gene point cloud.

#### 2.2.3 Knowledge Based MLP

In our adaptation of the traditional PointNet architecture, we have replaced the global max pooling layer with a knowledge-based one to incorporate biological pathway knowledge directly into the network. This innovative approach, inspired by the approach in [25], allows us to integrate specific gene pathway insights into the classification process. Initially, a convolutional MLP layer condenses each gene feature into a one-dimensional vector. Following this, we employ a mask linear transformation, utilizing the same methodology as described in [25], to weight features according to their biological relevance. The process culminates in a linear classifier that performs the final classification task based on the transformed features:

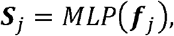

and

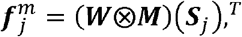

where ***S***_*j*_ ∈ R^4×60660^ represents the one-dimensional feature vector resultant from the convolutional MLP layer. Moreover, ***W***and ***M*** denote the linear transformation weights and the mask matrix, respectively.The operation ⨂ indicates element-wise multiplication across corresponding positions. The mask matrix ***M***, derived from GSEA [32], is a binary matrix encoding the presence or absence of genes within specific gene sets; each column correlates with a gene set, and each row corresponds to a gene. In ***M***, an entry *M*_*i,s*_ = 1sgnifies the inclusion of gene *i* in gene set *s*, and *M*_*i,s*_ = 0 indicates no association. This matrix is then replicated *m* times, matching the embedding dimension, to effectively transform gene inputs into gene set tokens, with m typically set to 48 but adjustable according to the needs of the analysis. The final classification output for sample *j*, 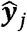, is obtained through:

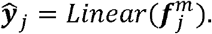

This knowledge-based MLP framework allows for a nuanced understanding of gene expression patterns in relation to known biological pathways, enhancing the model’s ability to identify relevant biomarkers and classify samples with a high degree of accuracy.

#### 2.2.4 Training Loss

As highlighted in Section 2.1, our training strategy incorporates both weighted cross-entropy loss and orthogonal loss tailored for the T-Net architecture to address the challenges posed by the unbalanced dataset. This approach ensures equitable learning across classes, particularly benefiting those with fewer samples by adjusting the convergence rate accordingly. The formulation of our loss function is as follows:

- The weighted cross-entropy loss, L_CE_, is calculated using:

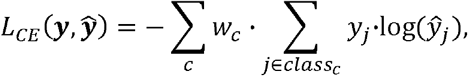

where ***y*** = [*y*_1_,*y*_2_, …,*y*_*C*_], and 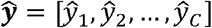 represent the ground truth and the predicted labels, respectively. The weight for each class *C, W*_*C*_, is derived as follows:

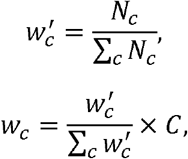

where *N*_*c*_ denotes the number of samples in class *c*. And *C* is the total class number.
- Orthogonal loss, designed to maintain the structural integrity of the transformations applied by the T-Net, is composed of two components:

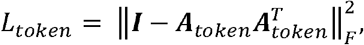

and

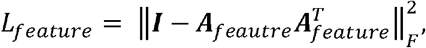

where ***A*** represents the alignment matrix produced by the T-Net, with ***A***_*token*_ and ***A***_*feautre*_ corresponding to the transformations for token and feature spaces, respectively, as visualized in **Figure 2(e)**. The total training loss, *L*_*total*_, combines these components, adjusting the influence of orthogonal loss via hyperparameters *λ*_1_ and *λ*_2_:

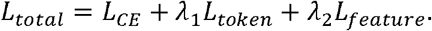

The hyperparameters *λ*_1_ and *λ*_2_ are tuned to balance the contribution of each loss component, optimizing the model’s performance across the diverse dataset. This comprehensive loss function framework enables our model to learn effectively from an unbalanced dataset, promoting the accurate classification of gene expressions while preserving the geometric properties encoded by the T-Net.

#### 2.2.5 Model Explanation

To transcend the traditional perception of deep learning models as inscrutable “black boxes,” we integrate inductive biases that mirror the underlying data structure, thereby bolstering the explainability of our model. This approach significantly enhances our ability to discern the model’s focal points, facilitating a deeper comprehension of genetic factors implicated in cancer pathogenesis. To achieve this, we employ a methodology akin to Gradient-weighted Class Activation Mapping (Grad-CAM) [19]. This technique illuminates the regions within the gene point cloud that the model deems crucial for its predictions, offering insights into the genetic landscape influencing cancer development. The attention score, 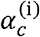 for gene *i* in class *c* is determined as follows:

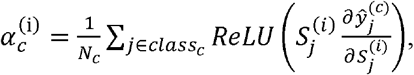

where 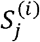 is the *i*th element of the one-dimensional feature vector ***S***_*j*_ as detailed in Section 2.2.3, and 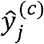 denotes the predicted probability that sample *j* belongs to class *c*.

In Section 4.4, we will delve into the top four genes that receive the most attention from the model. It is crucial to underline that the attention scores, 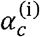 do not directly indicate whether the expression of these genes is increased or decreased in the context of class C cancer; rather, these scores highlight the genes’ relevance to the model’s classification decisions. This nuanced approach to model explanation not only enhances our understanding of the algorithm’s operation but also paves the way for more informed interpretations of the genetic underpinnings of cancer.

To mitigate the issue of overfitting, we employed a bootstrapping approach, conducting 10 iterations during the model testing phase. We randomly selected 60% of the data as the training set, ensuring that no category contained fewer than 10 data points; 20% was used as a validation set for determining hyperparameters and training cessation criteria. The remaining 20% of the data for each iteration served as the testing data. The composition of the test set was designed to mirror the overall dataset, maintaining a consistent proportion of data points across each category.

## 4. Results

In this section, we will discuss the performance of classification results, gene cluster results (the ability to push the related gene closer and verse vise), analysis of the related gene for each tumor class based on the attention score of the global gate pooling, and the ablation study. We run our model in V100-16GB GPU, without additional instructions, we set the hyperparameters as batch size is 8, learning rate is 0.0001, use step learning rate decay with step size is two and multiplicative factor is 0.5. We set *λ*_1_ = *λ*_1_ = 0.001.

### 4.1. Classification result

In this study, we explore the landscape of tumor classification by comparing various deep learning methodologies. Given the heterogeneity in network structures, we categorize these approaches into four distinct types and select a representative method from each for evaluation on our dataset. The absence of a universal benchmark dataset for tumor classification complicates direct comparisons across studies, as each employs different datasets for performance evaluation. Therefore, we reconstructed all the methods we selected, trained them on our dataset, and then tested them on the same cohort. The discernible performance disparities can offer insights into the efficacy of modeling gene expression data as point clouds.

Notably, TOSICA [25] is recognized as the state-of-the-art method for single-cell annotation. By juxtaposing its performance with our model’s, we aim to underscore our approach’s merits, particularly in the context of smaller datasets. For this comparison, we chose one model from each of the four network structure categories outlined in Section 2: CNN [22], ANN [8], SSAE [9], and TOSICA [25]. Due to the unavailability of code in the publications for CNN [22], ANN [8], and SSAE [9], we reconstructed these models based on the described architectures and hyperparameters. While TOSICA demonstrates superior performance on extensive single-cell datasets, our proposed GPNet exhibits a pronounced advantage on smaller datasets.

**Table 2** presents the classification performance metrics, where GPNet significantly outperforms the other models. This improvement underscores the suitability of representing gene expression data as point clouds, aligning closely with its intrinsic structure. Furthermore, incorporating inductive biases into the deep learning model enhances performance on smaller datasets. A statistical analysis confirms GPNet’s superiority over the second-best model, TOSICA [25], with a p-value of 0.00074 (indicating a confidence level greater than 99.9%).

**Table 2.**
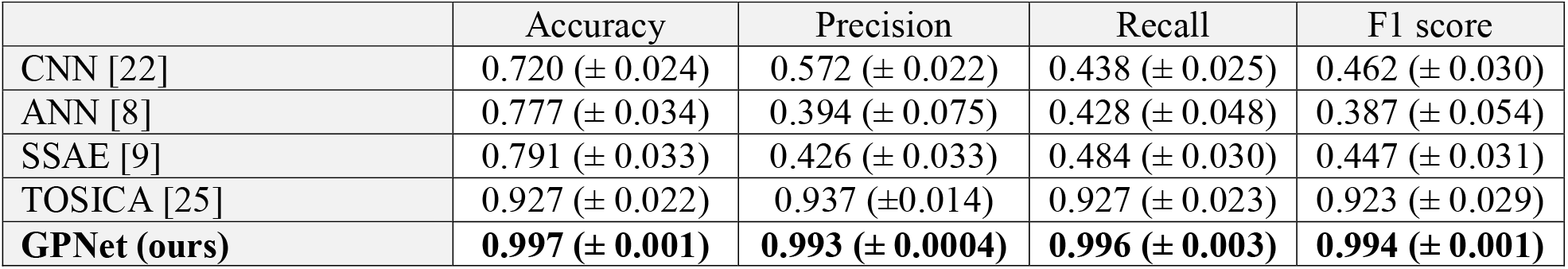
Classification result comparison for 6 tumor 10 classes’ datasets.

This comparative analysis not only validates GPNet’s conceptual premise but also highlights its practical superiority in leveraging the nuanced spatial relationships inherent in gene expression data for tumor classification.

#### Ablation study

**Table 3** presents an ablation study conducted for GPNet to evaluate the impact of various components on its performance. Specifically, we systematically removed the feature transform network, token transform network, and learnable normalization network to observe the effects on model efficacy. The results of this study reveal that omitting the token and feature transform networks does not significantly affect performance. This observation can be attributed to the consistent ordering of gene expression data inputs, which ensures that the “gene point cloud” remains inherently aligned. Consequently, the learnable alignment mechanisms provided by the token and feature transform networks do not contribute substantially in this context.

**Table 3.**
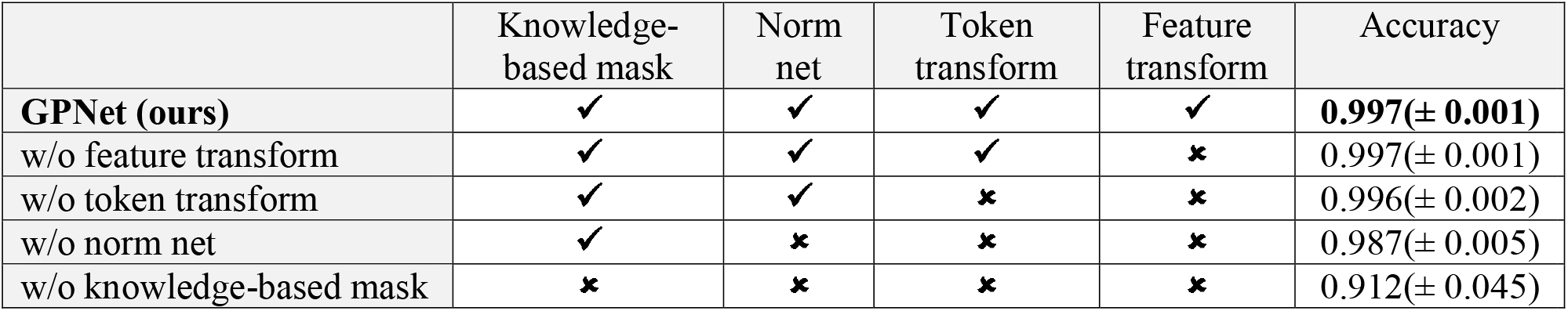
Ablation study for GPNet: Impact of Component Inclusion (⍰) and Exclusion (⍰) on Accuracy, where knowledge-based mask represents utilization of Knowledge-Based MLP, discussed in section 2.2.3, norm net represents utilization of Learnable Normalization Network, discussed in section 2.2.1, token transform represent the application of the first T-Net in PointNet for token transform, discussed in section 2.2.2, and Feature transform represent the application of the second T-Net in PointNet for feature transform, discussed in section 2.2.2.

|  | Knowledge-based mask | Norm net | Token transform | Feature transform | Accuracy |
| --- | --- | --- | --- | --- | --- |
| **GPNet (ours)** | ✓ | ✓ | ✓ | ✓ | **0.997(± 0.001)** |
| w/o feature transform | ✓ | ✓ | ✓ | ✗ | 0.997(± 0.001) |
| w/o token transform | ✓ | ✓ | ✗ | ✗ | 0.996(± 0.002) |
| w/o norm net | ✓ | ✗ | ✗ | ✗ | 0.987(± 0.005) |
| w/o knowledge-based mask | ✗ | ✗ | ✗ | ✗ | 0.912(± 0.045) |

Conversely, the learnable normalization network demonstrates a significant impact on model performance. This is likely due to the inherent characteristics of gene expression data, where normalization plays a critical role akin to that of Trimmed Mean of M-values (TMM) normalization in traditional gene expression analysis. The ablation study highlights the normalization network’ importance, showing a noticeable performance degradation in its absence.

### 4.2. Gene clustering result

We anticipated that the gene embedding layer would cluster related genes closely while separating unrelated ones. Following the approach of [33], we analyzed the learned gene relationships by extracting the gene point embeddings and applying unsupervised Leiden clustering, as shown in **Figure 3(a)**. This process resulted in 96 distinct clusters, with 35 of them containing more than 20 genes each, collectively encompassing over 95% of the total 60660 genes. We focused on these 35 clusters and queried the STRING database with the gene lists from each cluster to calculate network enrichment [34]. Notably, 74% (26 out of 35) of the gene clusters exhibited more interactions than expected by chance, as depicted in **Figure 3(b)**. This figure highlights the relationship between the expected and actual number of edges within gene networks, shedding light on the enrichment of these networks. Areas featuring densely clustered points, especially those of larger sizes and in purple, may indicate gene networks with strong enrichment signals, pinpointing critical areas for further investigation.

**Figure 3.**
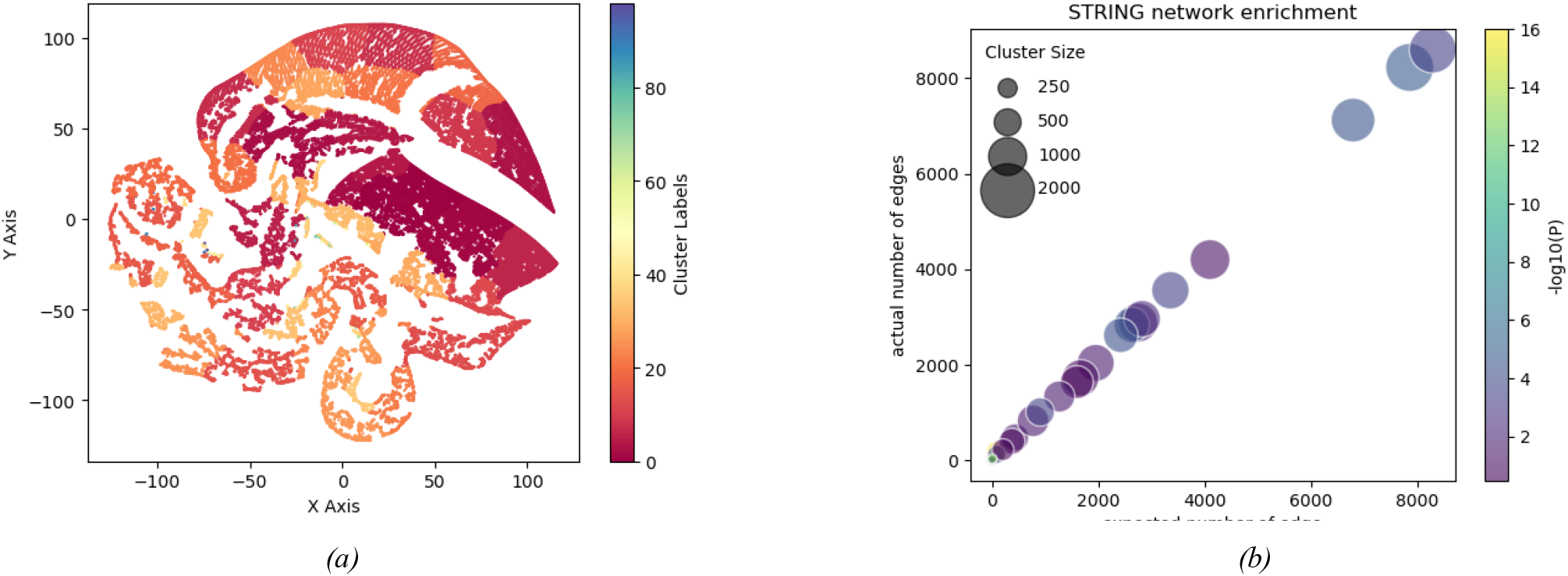
Learned gene point embedding similarity profiles reflect network biology. (a) t-SNE of global gene embedding colored by a different cluster. (b) STRING network enrichment plot of gene clusters.

### 4.3. Interpretation of models

In this subsection, we focus on identifying the top genes that received significant attention from the model during the classification process. To align our analysis with conventional differential expression analysis, we retrained the GPNet model for each specific type of cancer, contrasting it with its corresponding healthy tissue. Following the method for calculating attention scores outlined in Section 2.2.5, we determined an attention score for each gene. **Table 4** lists the top three genes that achieved the highest attention scores in distinguishing between tumor tissues and their corresponding normal tissues.

**Table 4.** Top 4 related genes to distinguish the tumor tissue and its corresponding normal tissue.

| Tumor type | Gene 1 | Gene 2 | Gene 3 |
| --- | --- | --- | --- |
| Acute Lymphoblastic Leukemia | LRMDA | PLA2G4A | SLC22A4 |
| Breast Invasive Carcinoma | MMP11 | VEGFD | COL10A1 |
| Colon Adenocarcinoma | OTOP2 | CA7 | BEST4 |
| Kidney Renal Papillary Cell Carcinoma | ATP6V1G3 | PRR35 | FRMD7 |
| Myeloid Leukemia | KCNA2 | BEND2 | MEOX1 |
| Plasma Cell Neoplasms | H4C13 | SEC11C | BHLHA15 |

By analyzing these genes, we discovered that certain genes, such as MMP11 in the context of breast invasive carcinoma, support the existing literature on their role in cancer progression [35, 36]. However, we did not find explicit studies regarding tumor development for some genes. For example, KCNA2 known as Kv1.2 voltage-gated potassium channel, encodes for the alpha subunit of the Kv1.2 voltage-gated potassium channel [37-39]. Although other members of this family of voltage-gated potassium channel subunits such as Kv2.1 have been indicated to play a role in the migration of prostate cancer as well as acting as potential reactive oxygen species sensors [40]. We did not find any explicit study to show KCNA2’s relationship with Myeloid Leukemia tumors. We will discuss the details of these in Section 5.

## Discussion

Our investigation into tumor classification through PGNet has yielded insights validating and extending current understandings within the field. The comparative analysis conducted in Section 4.1 illuminated the pivotal role of inductive biases in enhancing model accuracy. This was particularly evident in the distinctions drawn between PGNet and other models such as ANN and SSAE, where the deliberate incorporation of inductive biases significantly bolstered accuracy. A similar comparison with CNN models further emphasized the necessity of selecting appropriate inductive biases to optimize model performance. Additionally, the juxtaposition with the state-of-the-art TOSICA model underscored the unique advantage of PGNet in handling smaller datasets through the strategic introduction of inductive biases.

The experimental findings detailed in Section 4.2, where key components were sequentially omitted from PGNet, offered considerable insight into the model’s architecture. The negligible impact of removing the transform module on performance highlighted an intrinsic difference between gene profile data and conventional point cloud data, suggesting gene profile data’s inherent alignment, which obviates the need for transformation adjustments typically required for spatial data. Conversely, the observed enhancements attributed to the normalization module reaffirmed normalization’s critical role in gene profile analysis. Furthermore, the incremental benefits realized through integrating the knowledge-based MLP module underscored the value of embedding established gene information into the model, thereby leveraging inductive bias for improved performance.

In Section 4.3, our model’s ability to cluster similar genes into proximate spaces was demonstrated, validating PGNet’s capacity to accurately reflect functional gene relationships within its gene embedding space. This capability is crucial for understanding the complex interplay of genes in cancer pathogenesis and further supports the model’s conceptual framework.

Section 4.4 focused on PGNet’s interpretability, particularly through the lens of gene relevance in cancer occurrence. By spotlighting genes like MMP11 in the context of breast invasive carcinoma, we not only corroborate existing literature on its role in cancer progression but also highlight the model’s capability to identify genes with potential prognostic value. MMP11’s association with early breast cancer progression and its correlation with poor prognosis when expressed in stromal fibroblast-like cells adjacent to invasive ductal carcinoma exemplify how PGNet can illuminate critical biomarkers for cancer.

In examining specific genes highlighted by our model, we note the particular significance of MMP11, VEGFD, and COL10A1 in the context of breast invasive carcinoma. A temporary increase in MMP11 expression has been implicated as a potential early marker for breast cancer progression, especially before lymph node metastasis [35, 36]. The expression of MMP11 in stromal fibroblast-like cells adjacent to invasive ductal breast carcinoma has been correlated with critical prognostic factors, including tumor size and metastasis, pointing towards a poorer prognosis associated with higher MMP11 levels in both stromal and tumor cells [41].

VEGFD’s role, particularly in human invasive lobular breast cancer, highlights its contribution to promoting nodal metastasis, with significant expression in the cytoplasm of malignant cells and even within nuclei, indicating its active participation in invasive breast carcinomas [42]. VEGFC’s association as an independent indicator of poor prognosis further accentuates the complexity of angiogenic factors in cancer progression [43].

The analysis of COL10A1 - collagen type X alpha 1 chain, through bioinformatic approaches using The Cancer Genome Atlas (TCGA), revealed that COL10A1 mRNA was significantly overexpressed in multiple types of breast cancer [44]. Further analysis determined that overexpression of COL10A1 protein was linked to poor overall survival and advanced clinical stage, with the knockdown of COL10A1 significantly reducing breast cancer migration, invasion, and proliferation, as well as promoting apoptosis [34]. These findings were observed across various breast cancers, including invasive breast carcinoma, invasive ductal and lobular carcinomas [45].

However, not all genes identified by our model align with existing research across all cancer categories. For instance, in acute lymphoblastic leukemia (ALL), the top three genes (LRMDA, PLA2G4A, SLC22A4) lacked direct links to ALL but were strongly associated with other pathologies, including pancreatic ductal adenocarcinoma and myelogenous leukemia, among others. In kidney renal papillary cell carcinoma, ATP6V1G3 was directly linked to certain renal carcinomas such as KIRC, differentiating chromophobe RCC from other RCC types. Conversely, evidence for the association of lncRNA LINC02437 and PRR35 with kidney renal papillary cell carcinoma was minimal or non-existent, though related genes or gene paralogs have been linked to other cancers.

For myeloid leukemias, the top genes identified did not show a strong connection to the disease, despite substantial literature on related voltage-gated potassium channels and cancer proliferation. Similarly, none of the top genes identified for plasma cell neoplasms were directly linked to these conditions, although related snoRNA/scaRNA and lncRNAs have been associated with other cancers.

Our model’s attention mechanism, which differs from traditional differential expression analysis (DEA), focuses on the impact of changes in gene expression levels on classification outcomes rather than on traditional metrics such as p-values and fold changes. This approach uncovers genes that are crucial to the model’s decision-making process but may differ from those identified through DEA, thus necessitating traditional DEA for quantitative analysis.

Our model’s findings challenge conventional research paradigms by identifying genes with varying relevance across cancer types. For example, genes highlighted in acute lymphoblastic leukemia (ALL) and kidney renal papillary cell carcinoma demonstrate the model’s range, revealing associations with a broad spectrum of pathologies beyond the anticipated cancer types. This discrepancy highlights the complex relationship between genes and cancer, emphasizing the model’s refined approach to identifying potential biomarkers.

The use of an attention mechanism, distinct from conventional DEA, underscores the model’s novel approach, concentrating on the effects of gene expression changes on classification outcomes. This method provides essential insights into gene importance, but requires additional analyses for a thorough understanding.

## 5. Conclusions

This study advances the field of cancer classification by leveraging gene expression data to develop the Gene PointNet (GPNet) framework, a novel approach that treats RNA gene expression data as point clouds in high-dimensional space. By compiling a dataset from publicly available RNA-Seq gene expression data across four significant projects, Acute Myeloid Leukemia (AML), Breast Invasive Carcinoma (BRCA), Colon Adenocarcinoma (COAD), and Kidney Renal Papillary Cell Carcinoma (KIRP), our study encompasses 4916 samples from six different tumor types and 594 samples from adjacent normal cells, analyzing a total of 60660 genes per sample.

The GPNet framework is characterized by three main components: gene point cloud embedding, a lightweight PointNet backbone for feature extraction, and a knowledge-based MLP classifier that incorporates gene pathway knowledge for accurate cancer classification. This structure not only facilitates the efficient processing of gene expression data but also significantly enhances the model’s explainability with techniques such as grad-cam. Our approach, which models gene expression data as point clouds and incorporates inductive biases relevant to gene expression and pathways, has demonstrated substantial improvements in cancer classification accuracy, achieving an impressive accuracy rate of over 99%.

The innovative methodology of modeling gene expression data as high-dimensional point clouds via GPNet shows remarkable promise in the realm of cancer classification, particularly for small datasets. Integrating gene expression data with pathway knowledge underscores the vast potential of deep learning in oncology. Nonetheless, the challenge of ensuring model generalizability and addressing overfitting calls for further validations with external datasets. Future efforts will broaden the testing cohort to improve the model’s robustness and applicability across a wider array of cancer types. This endeavor will seek to validate GPNet’s utility in clinical settings and enhance its role in the discovery of biomarkers, thereby contributing significantly to the advancement of precision medicine in oncology.

## 6. Study Limitations

Our research, while pioneering in its approach to developing knowledge-based gene cloud representation for cancer classification using deep learning, has several limitations that warrant discussion and pave the way for future investigations.

### 1. Generalization and External Validation

One of the primary limitations of our study is the absence of external validation data. This gap restricts our ability to assess the model’s generalizability across different populations and clinical settings. The lack of external validation data underscores the necessity of conducting further research to evaluate the model’s findings across a broader range of cohorts. This will be crucial for confirming the model’s clinical applicability and ensuring that it can reliably support medical decisions in diverse healthcare environments.

### 2. Dataset Imbalance and Overfitting

Although we have employed strategic adjustments and bootstrapping strategies to mitigate the risks of dataset imbalance and overfitting, these challenges remain significant concerns in machine learning applications within genomics. Ensuring the model’s robustness and accuracy in the face of imbalanced data requires continuous refinement of our methodological approaches.

### 3. Scope of Gene Relevance Identification

Our model’s attention mechanism offers a novel perspective on identifying genes relevant to cancer classification. However, this approach may diverge from traditional differential expression analysis (DEA), highlighting genes based on their impact on classification outcomes rather than their differential expression levels. This discrepancy necessitates using traditional DEA for comprehensive gene expression studies, potentially limiting the scope of biomarker identification using our model alone.

### 4. Interpretability and Biological Insights

While our model provides valuable insights into gene roles in cancer pathology, the complexity of biological systems and the nuanced interplay between genes and cancer pathology may limit the interpretability of the model’s findings. Bridging computational findings with deeper biological understanding remains challenging, requiring collaborative efforts between computational scientists and biologists.

In addressing these limitations, future work will focus on obtaining external validation datasets, enhancing methodological approaches to tackle dataset imbalance, and fostering interdisciplinary collaborations. These efforts will aim to reinforce the model’s predictive capabilities, broaden its applicability, and contribute to advancing precision oncology.

## 7. Acknowledgement

This work has already submitted to Neural Computing and Applications. This effort sponsored by the U.S. Government under HDTRA 12310003, “Host signaling mechanisms contributing to endothelial damage in hemorrhagic fever virus infection,” PI: Narayanan. The US Government is authorized to reproduce and distribute reprints for Governmental purposes, notwithstanding any copyright notation thereon. The views and conclusions contained herein are those of the authors and should not be interpreted as necessarily representing the official policies or endorsements, either expressed or implied, of the U.S. Government.

